# Genetic diversity and antimicrobial resistance of *Campylobacter jejuni* isolates from Gambian children under five with moderate-to-severe diarrhoea and healthy Controls

**DOI:** 10.1101/2024.07.30.605890

**Authors:** Modupeh Betts, Michel Dione, Usman Nurudeen Ikumapayi, Madikay Senghore, Modou Lamin, Ebenezer Foster-Nyarko, James Jaffali, Sandra Panchalingam, James P Nataro, Karen L Kotloff, Myron M Levine, Debasish Saha, Jahangir Hossain, Brenda Kwambana-Adams, Martin Antonio

## Abstract

**Introduction:** *Campylobacter* is a leading cause of bacterial gastroenteritis globally, but its molecular epidemiology remains poorly understood in sub-Saharan Africa. This study investigates the genotypic population structure of *Campylobacter jejuni* isolates from children with moderate-to-severe diarrhoea (MSD) and healthy controls in The Gambia. Additionally, we determined the antimicrobial susceptibility levels of the isolates.

**Methods:** As part of the Global Enteric Multicenter Study (GEMS) in The Gambia, a total of 49 *C. jejuni* isolates were collected from the stools of children under 5 years old, including 22 with MSD and 27 healthy controls. These isolates were subjected to multilocus sequence typing (MLST) and antimicrobial susceptibility testing using the disc-diffusion method.

**Results:** The *C. jejuni* isolates belonged to 22 sequence types (STs), ten of which were novel. The most common STs were ST-353 (19.1%, 9/47), ST-7784 (12.7%, 6/47), and ST-1038 (10.6%, 5/47). All isolates were fully susceptible to erythromycin, tetracycline, gentamicin and chloramphenicol, with two isolates (4.4%, 2/45) resistant to ciprofloxacin and nalidixic acid. Antimicrobial resistance or intermediate susceptibility to trimethoprim-sulfamethoxazole, cefotaxime and ampicillin was observed in 91.1% (41/45), 90.9% (40/44), and 44.4% (20/45) of the isolates, respectively. There was no strong evidence linking *C. jejuni* antimicrobial susceptibility or MLST genotype to MSD status.

**Conclusion:** This study provides the first overview of the high genotypic diversity of human *C. jejuni* isolates in The Gambia and reveals low-level resistance among the isolates to antibiotics commonly used to treat campylobacteriosis. The study contributes to understanding the epidemiology and resistance patterns of *C. jejuni* in this region.

## Introduction

Campylobacter infections (campylobacteriosis) are a leading cause of bacterial gastroenteritis globally, accounting for approximately 14% of all diarrhoeal cases (Kaakoush *et al*., 2015; Kirk *et al*., 2015; Sherman *et al.,* 2010). In developing countries, *Campylobacter* is the most frequently isolated bacterial pathogen among paediatric cases of diarrhoea (Coker *et al*. 2002, Platts-Mills *et al*. 2015). At least 90% of campylobacteriosis are due to the subspecies *Campylobacter jejuni* (Allos, 2001). Most human campylobacteriosis cases are self-limiting and only require supportive treatment (Allos 2001, Snelling *et al*. 2005). However, infections in immunocompromised individuals, patients with prolonged diarrhoea, and cases of septicaemia require antibiotic therapy (Allos 2001, Butzler 2004, Moore *et al*. 2006). Antibiotic treatment may also be crucial for preventing potential post-infectious sequelae such as Guillain-Barré syndrome, irritable bowel syndrome, and reactive arthritis (Halvorson *et al*. 2006, WHO 2013). The antibiotics of choice for treating campylobacteriosis are macrolides and fluoroquinolones (Kaakoush *et al*. 2015). However, increasing antibiotic resistance in *Campylobacter* strains from both humans and animals has become a significant global health challenge. The World Health Organization has listed *Campylobacter* as a high-priority antibiotic-resistant bacterium (WHO 2017, Tacconelli *et al*. 2018). Consumption and handling of contaminated poultry and poultry products are the most common risk factors associated with *C. jejuni* infection (Kaakoush *et al*. 2015).

Data on the molecular epidemiology and genetic diversity of *Campylobacter* is not available for The Gambia and is sparse for the rest of sub-Saharan Africa (WHO 2013, Kaakoush *et al*. 2015). As early as 1979, *Campylobacter* infection was identified as an important cause of gastroenteritis among Gambian children under 5 years old, accounting for about twice as many cases as *Shigella* and *Salmonella* (Billingham 1981). The Global Enteric Multicenter Study (GEMS) was conducted in The Gambia and six other low-and-middle-income countries in Sub-Saharan Africa and South Asia from 2007 to 2010 (Kotloff *et al*. 2013). Findings from GEMS showed significant associations between *C jejuni* infections and diarrhoeal disease in children under 5 years in the three Asian sites (India, Bangladesh, Pakistan), but not in the sub-Saharan African sites (The Gambia, Kenya, Mozambique, Mali) (Kotloff *et al*. 2013). In The Gambia, the proportion of *C. jejuni* isolated from stools of moderate-to-severe diarrhoea (MSD) cases and healthy controls was identical (Kotloff *et al*. 2013).

We investigated the genotypic structure of *C. jejuni* in The Gambia and explored the hypothesis that *C. jejuni* isolated from MSD cases and controls have distinct genotypes. We used Multilocus sequence typing (MLST) to characterise *C. jejuni* isolates from The Gambia during GEMS and determined antimicrobial susceptibility patterns using disk diffusion method. MLST is renowned for its capacity to decipher the genetic epidemiology of bacterial pathogens and allows for comparisons of the population structure of strains from different locations (Dingle *et al*. 2002, Sails *et al*. 2003).

This study contributes a baseline understanding of the population dynamics among *C. jejuni* isolates in The Gambia, provides an update on antibiotic susceptibility patterns of the isolates, and offers insights into genetic relatedness to circulating strains from other parts of sub-Saharan Africa.

## Methods

### GEMS study setting in The Gambia

The Global Enteric Multicenter Study (GEMS) was a three-year, prospective, age-stratified, matched case-control study of moderate-to-severe diarrhoea (MSD) conducted in the Upper River Region of The Gambia from December 2007 to December 2010. The study focused on children aged 0-59 months residing in a population under demographic surveillance system (DSS) (Kotloff *et al.,* 2013). Stool samples were collected from children presenting with MSD at healthcare facilities covered by the DSS. For each MSD case, stool samples were also collected from 1-3 diarrhoea-free controls matched by age, sex, and residence. Subjects were recruited into three age strata: 0-11 months, 12-23 months, and 24-59 months. Detailed descriptions of case definitions, inclusion criteria, and participant recruitment have been previously published (Kotloff *et al.,* 2012).

### Ethics

Study participants were enrolled, and stool samples collected only after obtaining written informed consent from their parents or guardians. Ethical approval for the study was granted by the joint Medical Research Council Unit The Gambia at the London School of Hygiene and Tropical Medicine and the Government of The Gambia ethics committee.

### *Campylobacter* isolation and antimicrobial susceptibility testing

The methods for identifying *Campylobacter* in GEMS were detailed by Panchalingam *et al*. (2012). Pure colonies of *C. jejuni* were suspended in 2 mL of tryptone soya broth (TSB) to achieve turbidity equivalent to a 0.5 McFarland standard. The suspension was then inoculated and uniformly spread onto Mueller-Hinton agar supplemented with 5% sheep blood using sterile swabs (Sterilin, UK). The Kirby-Bauer disc diffusion technique was employed to test susceptibility to nine antimicrobials (Oxoid, UK): ciprofloxacin (5 µg), nalidixic acid (30 µg), erythromycin (15 µg), tetracycline (30 µg), ampicillin (10 µg), gentamicin (10 µg), cefotaxime (30 µg), trimethoprim-sulfamethoxazole (1.25/23.75 µg), and chloramphenicol (30 µg). Plates were incubated at 42°C under microaerophilic conditions for 48 hours. Susceptibility was determined by measuring the zone of inhibition in millimetres and interpreting results according to Clinical and Laboratory Standards Institute (CLSI) guidelines for Enterobacteriaceae (CLSI, 2014).

### DNA extraction and multilocus sequence typing (MLST)

Genomic DNA was extracted from liquid cultures of pure *C. jejuni* colonies grown in TSB using the automated NucliSens® easyMAG™ nucleic acid extraction system (Biomerieux, France). A 100 µL aliquot of the *C. jejuni* suspension was added to 2 mL of lysis buffer, vortexed for 1 minute to achieve homogeneity, and incubated overnight at 4°C. The NucliSens® easyMAG™ generic protocol was used with a final elution volume of 100 µL. MLST PCR and sequencing reactions were performed using primers and conditions for seven housekeeping genes (*aspA, glnA, gltA, glyA, pgm, tkt,* and *uncA*) as previously described (Dingle *et al.,* 2001). Sequencing of the housekeeping gene amplicons was conducted at Macrogen Inc., South Korea.

### Bioinformatics analyses

Sequences were aligned and assembled using LaserGene DNA Star (v7.1) software. Alleles, sequence types (STs), and clonal complexes (CC) were identified by querying the MLST website from the *Campylobacter* database (https://pubmlst.org/campylobacter/). Novel alleles and ST profiles were submitted to the database curator for assignment of new alleles or STs. Clustering and minimum spanning tree construction were performed using BioNumerics (v6.6) software. Maximum likelihood (ML) phylogeny was reconstructed based on the concatenated MLST alleles using the General Time Reversible model with 100 replicates in SeaView (v4).

## Results

### Prevalence of *Campylobacter* in children with MSD and healthy controls

A total of 2,598 children under five years old were enrolled in the GEMS study in The Gambia, comprising 1,029 MSD cases and 1,569 healthy controls (Kotloff *et al*., 2012). *Campylobacter* species were isolated from 105 independent stool samples, yielding an overall prevalence of 4.0%, with similar rates between MSD cases (4.0%, 41/1,029) and healthy controls (4.1%, 64/1,569) (**Table 1**). The majority of *Campylobacter* species (89.5%, 94/105) were isolated from children under two years old. Approximately 57% (60/105) of the *Campylobacter* isolates were classified as *C. jejuni*, which are the focus of this study (**Table 1**). Due to missing or non-viable isolates, MLST and antimicrobial susceptibility testing were conducted on the remaining 49 (81.7%) *C. jejuni* isolates.

**Table 1:**
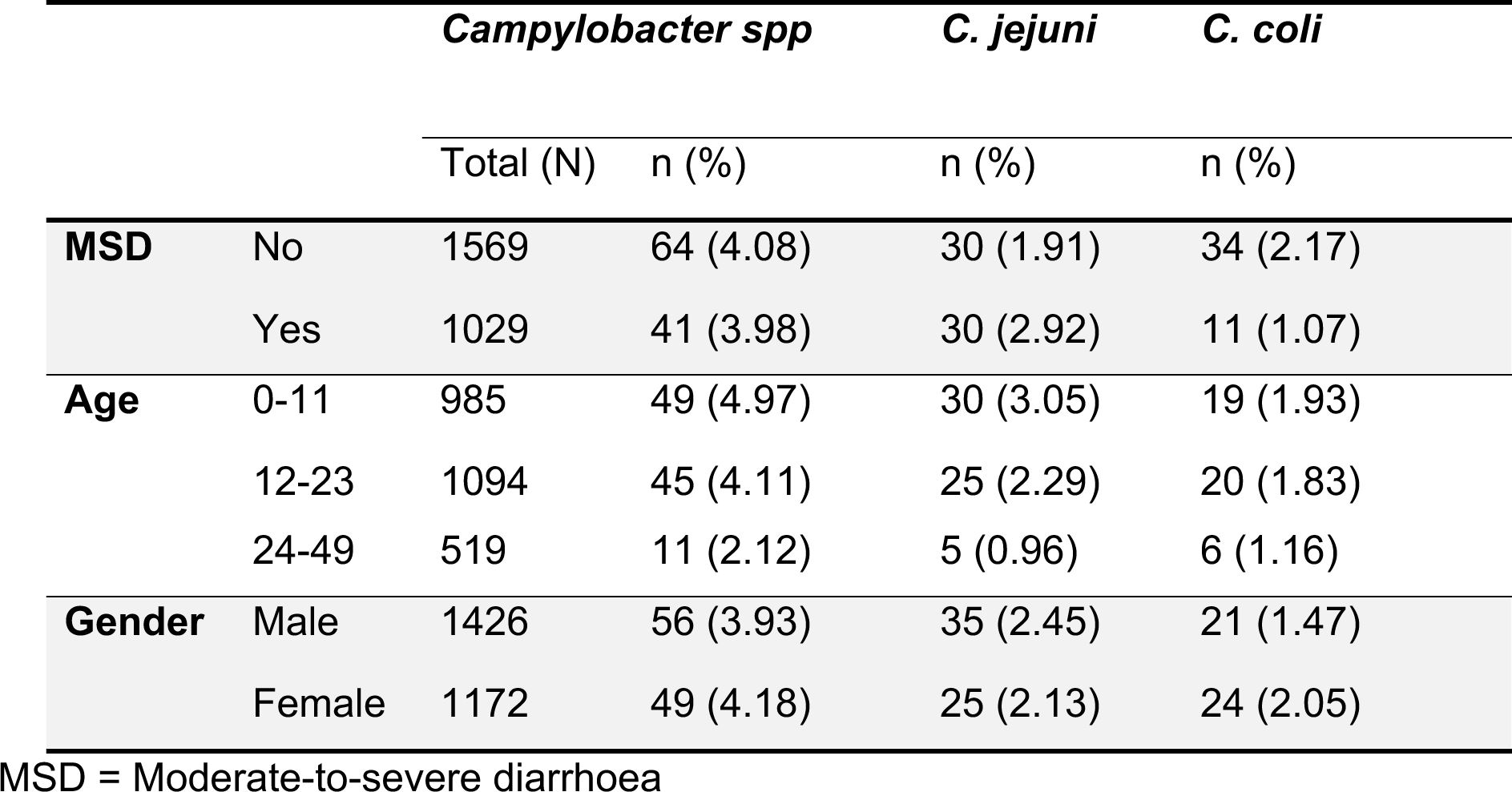
Distribution of *Campylobacter* species isolated from the stools of children five years old during the GEMS study in The Gambia (December 2007 – December 2010)

### Genetic diversity of *C. jejuni* isolates in Gambian children

Complete MLST profiles were obtained for 47 of the 49 (95.9%) *C. jejuni* isolates, revealing 22 different STs and indicating high genetic diversity. Ten novel STs (ST-7772, ST-7782, ST-7783, ST-7784, ST-7790, ST-7791, ST-7792, ST-7793, ST-8112, and ST-8113) and one novel allele for the *tkt* loci (assigned allele 491) were identified (**Supplementary Table 1**). The *tkt* 491 allele has a single nucleotide difference to *tkt* allele 3. The most common genotypes were ST-353 (19.1%, 9/47), the novel ST-7784 (12.8%, 6/47), ST-1038 (10.6%, 5/47), ST-607 (8.5%, 4/47), and ST-52 (8.5%, 4/47). ST-1036 and ST-1039 were each found in 2/47 (4.3%) isolates, while fifteen other STs were each detected once. Overall, the novel STs comprised 31.9% (15/47) of the total isolates.

A total of 43/47 (91.5%) isolates were distributed into 10 clonal complexes (CCs), with CC353 (42.6%, 20/47), CC607 (14.9%, 7/47), CC354 (12.8%, 6/47), and CC52 (8.5%, 4/47) being the most prevalent. Other clonal complexes identified included CC22, CC206, CC362, CC460, CC48, and CC574, each represented by a single isolate. Three STs (ST-1039, ST-7772, and ST-7793) did not belong to any clonal complex (**Supplementary Table 1**).

### MLST genotype distribution in MSD cases vs healthy controls

The two most prevalent STs, ST-353 and ST-7784, were found equally in isolates from both MSD cases and healthy controls (**Figure 1**). However, certain MLST genotypes were more common in one group compared to the other. For instance, 80% (4/5) of isolates with the ST-1038 genotype were from MSD cases, whereas 75% (3/4) of isolates with the ST-607 genotype were from healthy controls (**Figure 1**). The small sample size limits the power of this study to draw definitive conclusions about these associations, necessitating further investigation.

**Figure 1:**
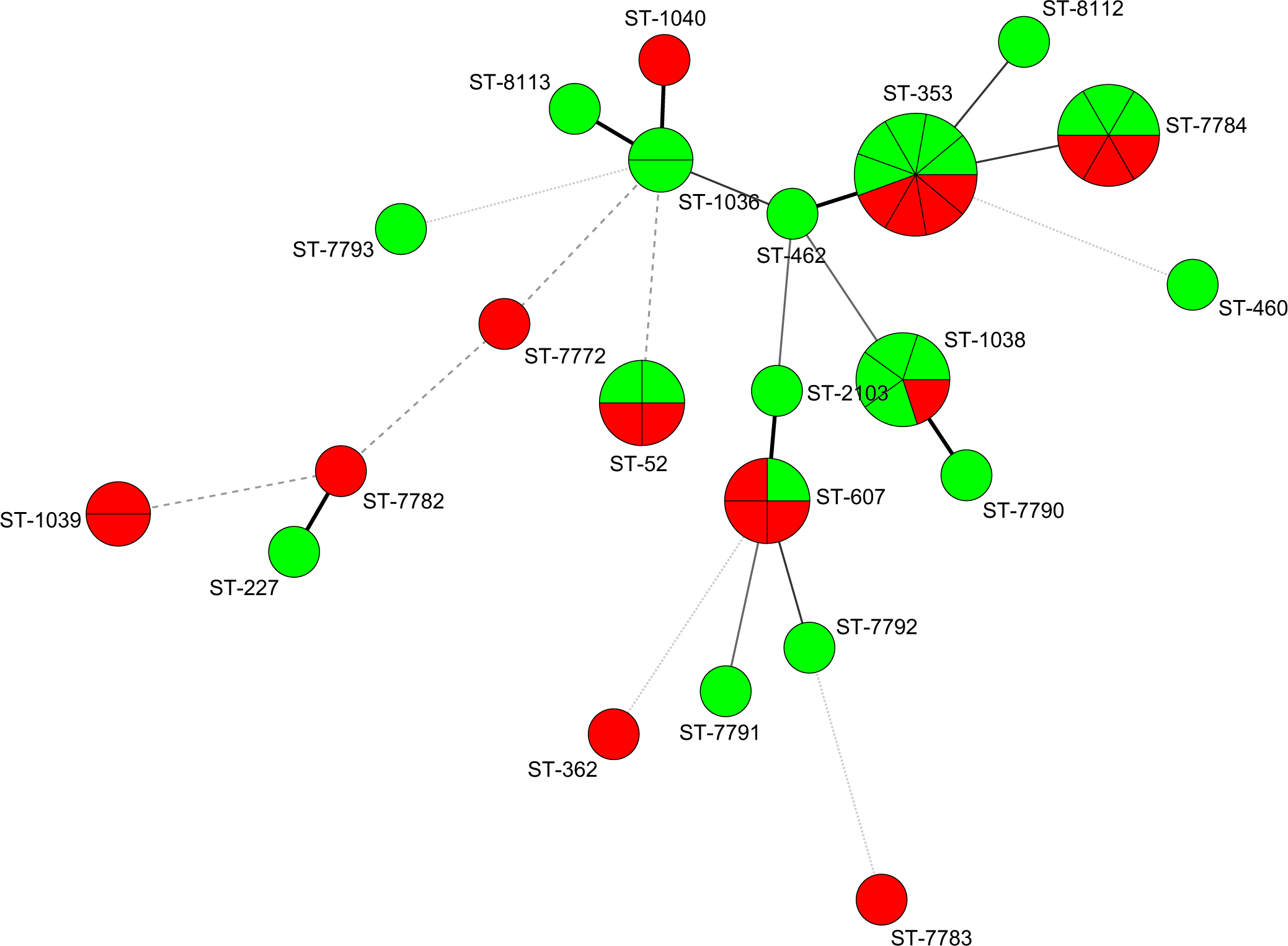
Clustering of MLST genotypes of *C. jejuni* from stools of children with moderate-to-severe diarrhoea (MSD) and healthy controls using minimum spanning tree. Each circle represents an ST profile, with the area of each circle corresponding to the number of isolates with that profile. The length of the lines represents the number of locus variants: Thick, short, solid lines connect single-locus variants; thick longer solid lines connect double-locus variants; thin, long solid lines connect triple-locus variants; dashed lines connect quadruple-locus variants, and dotted lines connect quintuple-locus variants. Red segments represent MSD cases, and green segments represent healthy controls.

### Antimicrobial susceptibility of *C. jejuni* isolates from Gambia children

Antimicrobial susceptibility data were available for 45 of the 49 (91.8%) *C. jejuni* isolates. For 4 out of 49 (8.2%) isolates for which MLST genotypes were obtained, antimicrobial susceptibility data was not available. All isolates tested (n=45) were fully susceptible to erythromycin, tetracycline, chloramphenicol, and gentamicin (**Table 2**). Susceptibility to trimethoprim-sulfamethoxazole, cefotaxime, and ampicillin was observed in 8.9% (4/45), 9.1% (4/44), and 55.6% (25/45) of the isolates, respectively. The fluoroquinolones ciprofloxacin and nalidixic acid had identical profiles, with 95.6% (43/45) of isolates susceptible and 4.4% (2/45) resistant to both agents. One isolate from an MSD case with the ST-1039 genotype was susceptible to all nine antimicrobials tested. The antimicrobial profile for the other ST-1039 isolate was unavailable, leaving it undetermined whether pan-susceptibility is characteristic of this genotype (**Figure 2**). Approximately 37.8% (17/45) of isolates exhibited resistance or intermediate susceptibility to three or more antimicrobials, suggesting the presence of potentially multi-drug resistant (MDR) strains (**Figure 2**). Disk diffusion method-generated antimicrobial susceptibility profiles will need confirmation via the minimum inhibitory concentration (MIC) by E-test or agar dilution method to verify MDR profiles.

**Figure 2:**
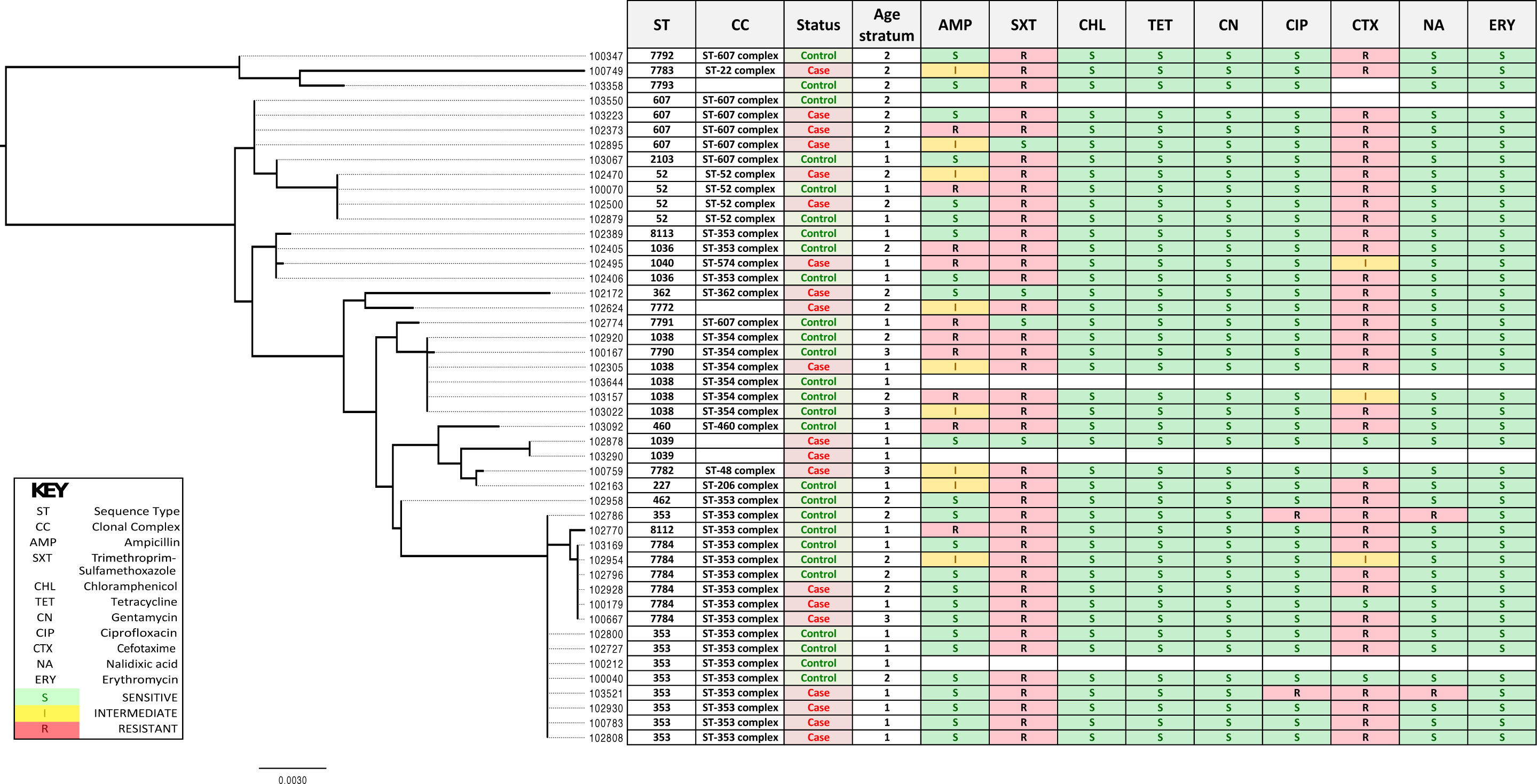
Maximum likelihood phylogeny of 7 concatenated MLST alleles. The phylogeny is presented alongside participant metadata and isolate antimicrobial susceptibility profiles. Age strata are indicated as follows: 1= 0-11 months, 2= 12-23 months, 3= 24-59 months.

**Table 2:** Susceptibility of *C. jejuni* isolates from Gambian children under five to nine antimicrobials agents.

| Antimicrobial agent (concentration) | *C. jejuni* isolates (n) |  |  |  |
| --- | --- | --- | --- | --- |
|  | S | I | R | S (%) |
| Ampicillin (10µg) | 25 | 10 | 10 | 55.6 |
| Trimethoprim-Sulfamethoxazole (25µg) | 4 | 0 | 41 | 8.9 |
| Chloramphenicol (30µg) | 45 | 0 | 0 | 100 |
| Tetracycline (30µg) | 45 | 0 | 0 | 100 |
| Gentamicin (10µg) | 45 | 0 | 0 | 100 |
| Ciprofloxacin (5µg) | 43 | 0 | 2 | 95.6 |
| Cefotaxime (30µg) | 4 | 3 | 37 | 90.9 |
| Nalidixic acid (30µg) | 43 | 0 | 2 | 95.6 |
| Erythromycin (15µg) | 45 | 0 | 0 | 100 |

S = Susceptible; I = Intermediately susceptible; R = Resistant

### Relationship between antimicrobial susceptibility and MLST genotypes

Complete antimicrobial susceptibility and MLST genotype data were obtained for 43 of the 49 (87.8%) *C. jejuni* isolates. A maximum likelihood (ML) phylogeny based on the concatenated MLST sequences of each isolate was reconstructed and compared to the respective antimicrobial profiles (**Figure 2**). No clear correlation was observed between antimicrobial susceptibility patterns and MLST genotypes or MSD status in this dataset. Isolates with the same MLST genotype clustered together in the ML phylogeny often exhibited different antimicrobial profiles, regardless of MSD status or age strata (**Figure 2**). The two fluoroquinolone-resistant isolates (one from an MSD case aged 0-11 months and the other from a healthy control aged 12-23 months) exhibited a unique antimicrobial resistance profile, being resistant to trimethoprim-sulfamethoxazole, cefotaxime, ciprofloxacin, and nalidixic acid. These isolates shared the same MLST profile (ST-353). However, other ST-353 isolates were susceptible to both ciprofloxacin and nalidixic acid, indicating variability in antimicrobial resistance within the same genotype (**Figure 2**).

## Discussion

In this study, we used multilocus sequence typing (MLST) to describe for the first time the circulating genotypes of *Campylobacter jejuni* isolates obtained from children under five years old with moderate-to-severe diarrhoea (MSD) and healthy controls in The Gambia. We also evaluated the susceptibility of these isolates to commonly used antibiotics.

Globally, antibiotic resistance in *Campylobacter* species has been increasing. Consisted with the findings by Billingham (1981) from four decade ago, our study found that all *C. jejuni* isolates in The Gambia were fully susceptible to erythromycin, tetracycline, chloramphenicol, and gentamicin. The antibiotics of choice for the treatment of campylobacteriosis are macrolides (e.g., erythromycin, azithromycin) and fluroquinolones (e.g., ciprofloxacin, nalidixic acid) (Kaakoush *et al*. 2015). However, we identified fluoroquinolone-resistant *C. jejuni* strains, with 4.4% (2/45) of isolates resistant to both ciprofloxacin and nalidixic acid. Furthermore, high levels of resistance were observed against ampicillin, trimethoprim-sulfamethoxazole, and cefotaxime. These antibiotics were not tested in the earlier study by Billingham (1981), so we cannot definitively conclude if resistance has increased over time. The association between antibiotic use in food animals and the rise of antimicrobial-resistant *Campylobacter* strains in humans is well-documented (Endtz 1991, Allos 2001, McCrackin *et al*. 2016, Hoelzer *et al*. 2017). However, data on antimicrobial usage in Gambian food animals is limited, but likely lower than in developed countries. Additionally, confirmation of resistant strains using alternative methods, such as the minimum inhibitory concentration (MIC) method, was not performed, which might have affected our observed resistance frequency (Lehtopolku et al. 2012).

Our study revealed high genetic diversity among the *C. jejuni* strains, identifying 22 sequence types (STs) and 10 clonal complexes (CCs) from 47 isolates. Notably, some common human genotypes, such as CC21 and CC45, were absent (Dingle *et al*. 2001, Dingle, Colles *et al*. 2002, Schouls *et al*. 2003, Levesque *et al*. 2008, de Haan *et al*. 2010). The predominant clone in The Gambia was CC353, prevalent in West Africa and associated with both human and animal sources (Kinana *et al*. 2006, Ngulukun *et al*. 2016). In our study, CC353 was present in both MSD cases and healthy controls, conflicting with findings from Europe where CC353 was linked to disease in children (Ramonaite *et al*. 2014). Other common clones in our data, such as CC607, CC354, and CC52, which are also associated with disease in Europe, were identified in both MSD cases and healthy controls (**Supplement Table 1**). The second most common genotype, the novel ST-7784, belonged to the same clonal complex (CC353) as the prevalent ST-353, indicating genetic divergence within this complex (**Figure 2**).

Although certain STs, such as ST-607 and ST-1038, were more commonly identified in MSD cases or controls, our limited sample size prevents definitive conclusions about the association between MLST genotypes and MSD status (**Figure 1**). Previous studies have shown mixed results regarding the correlation between MLST genotypes and antibitic resistance patterns (Kinana *et al*. 2006, Levesque *et al*. 2008, Shin *et al*. 2013, Cha *et al*. 2016). Our findings align with studies by Kinana *et al*. (2006) and Levesque *et al*. (2008), which found no clear association between antibiotic resistance profiles and *C. jejuni* genotypes.

The present study has some limitations. First, the culture method used in GEMS has been shown to under-detect *Campylobacter spp* by twice compared to quantitative molecular diagnostic approaches (Liu *et al*. 2016), potentially missing other circulating genotypes. For example, the ST-2928 genotype, which was first described from a child resident in the urban area of The Gambia and belonging to CC443, was not present in our study (Morris *et al*. 2008). Second, the MLST methodology indexes sequence variation in only seven housekeeping genes, whereas whole genome sequencing (WGS) could provide higher discriminatory power and identify antibiotic resistance genes. Third, the small sample size limits the statistical power to test associations between genotypes and disease status and to draw definitive conclusions. Finally, the MSD cases and controls in this sub-study were not matched.

Contrary to the early report by Billingham (1981), *Campylobacter* was not a major cause of diarrhoea during GEMS in The Gambia (Kotloff *et al*. 2013). The findings from GEMS were more aligned with a longitudinal study in The Gambia showing higher isolation of *Campylobacter* during control periods than diarrhoeal episodes (Rowland *et al*. 1985). Despite this, *Campylobacter* infections are crucial for child health, as both symptomatic and asymptomatic infections are associated with growth faltering in developing countries (Lee *et al*. 2013, Amour *et al*. 2016). Consistent with the epidemiology of *Campylobacter* in developing countries, *Campylobacter* was predominantly detected from the stools of children 0-23 months old in GEMS.

Our study highlights the high genotypic diversity of *C. jejuni* and identifies ten novel STs, underlining the unique population structure in The Gambia. The high susceptibility to erythromycin and alternative antibiotics supports their continued use for treating campylobacteriosis. Given the scarcity of data from this region, our study contributes to an important knowledge gap in the epidemiology of *C. jejuni* infections in Africa and may inform future vaccine development strategies (Monteiro *et al*. 2009, Nothaft *et al*. 2016). Future research should include larger sample sizes and samples from animals and environmental sources to provide a more comprehensive understanding of *C. jejuni* epidemiology in The Gambia.

## Acknowledgement

The GEMS study was supported by the Bill and Melinda Gates Foundation through the University of Maryland School of Medicine, Baltimore, USA. This sub-study was funded and facilitated by MRC Unit The Gambia at the London School of Hygiene and Tropical Medicine. We extend our gratitude to the dedicated field, clinical, and laboratory teams whose efforts made this research possible. We thank the study participants and their parents/guardians for their invaluable contribution to GEMS.

---

**Supplementary Table 1:**
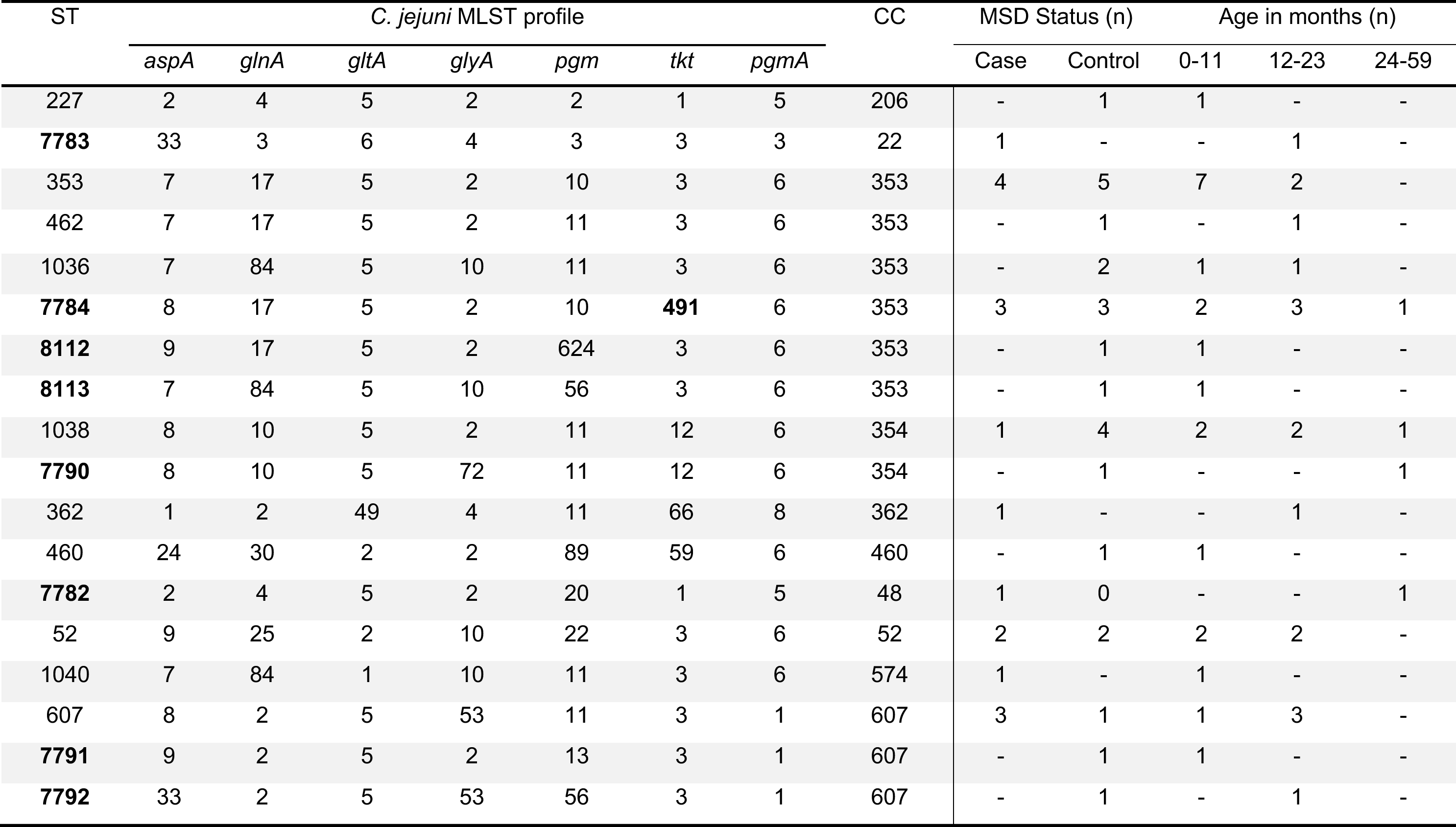
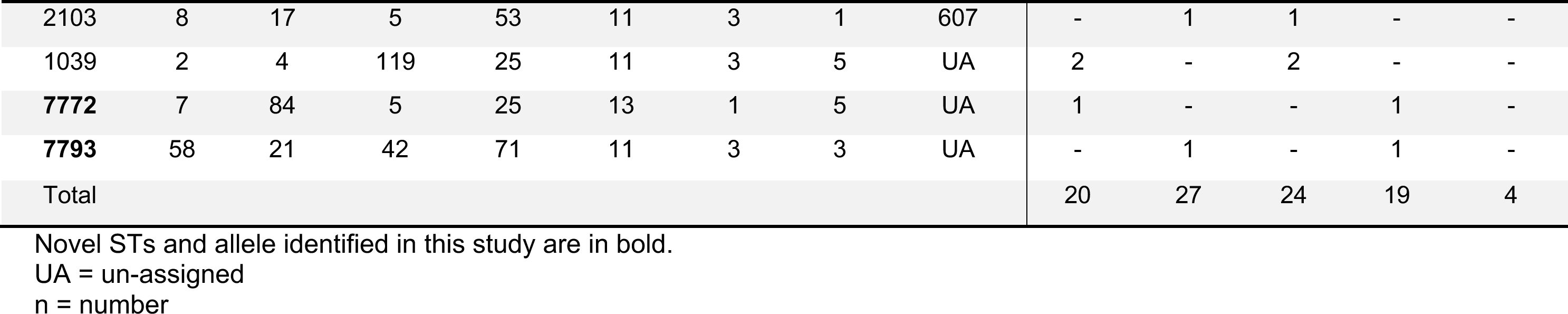
Distribution of *Campylobacter jejuni* MLST genotypes across three age strata in Gambian children under 5 years old with moderate-to-severe diarrhoea (MSD) and healthy controls.

